# Molecular mechanism of symmetry breaking in a 3D model of a human epiblast

**DOI:** 10.1101/330704

**Authors:** Mijo Simunovic, Jakob J. Metzger, Fred Etoc, Anna Yoney, Albert Ruzo, Iain Martyn, Gist Croft, Ali H. Brivanlou, Eric D. Siggia

**Affiliations:** Laboratory of Stem Cell biology and Molecular Embryology, The Rockefeller University, New York, NY 10065, USA; Center for Studies in Physics and Biology, The Rockefeller University, New York, NY 10065, USA

## Abstract

Breaking the anterior-posterior (AP) symmetry in mammals takes place at gastrulation. Much of the signaling network underlying this process has been elucidated in the mouse, however there is no direct molecular evidence of events driving axis formation in humans. Here, we use human embryonic stem cells to generate an *in vitro* 3D model of a human epiblast whose size, cell polarity, and gene expression are similar to a 10-day human epiblast. A defined dose of bone morphogenetic protein 4 (BMP4) spontaneously breaks axial symmetry, and induces markers of the primitive streak and epithelial to mesenchymal transition. By gene knockouts and live-cell imaging we show that, downstream of BMP4, WNT3 and its inhibitor DKK1 play key roles in this process. Our work demonstrates that a model human epiblast can break axial symmetry despite no asymmetry in the initial signal and in the absence of extraembryonic tissues or maternal cues. Our 3D model opens routes to capturing molecular events underlying axial symmetry breaking phenomena, which have largely been unexplored in model human systems.

## INTRODUCTION

Gastrulation initiates the morphogenetic movements that establish the vertebrate body plan by creating the three body axes: anterior-posterior (AP), dorsal-ventral (DV) and the left-right axes^1, 2^. Just prior to gastrulation, the mammalian embryo consists of a polarized epithelium, the epiblast, which will give rise to all the cells of the fetus proper. The epiblast is enveloped in various extra-embryonic tissues that supply essential developmental signals and nutrients^3^. While in lower vertebrates, such as amphibians, the asymmetry in RNA and protein distribution in the egg coupled with the location of sperm entry orients the AP and DV axes; in mammals, the choice of axis direction appears random^4^. Hence the question: how do the embryonic and extraembryonic signaling pathways cooperate to ensure unique body coordinates?

At the onset of gastrulation in the mouse, WNT and NODAL initiate the primitive streak in the posterior axis, and their inhibition leads to specification of the anterior fates. In the extraembryonic tissues the posterior visceral endoderm expresses WNT3, which is required for the proper timing of the primitive streak formation^5^. One of key sources of inhibitors of WNT and NODAL in the mouse is the anterior visceral endoderm (AVE), whose failure to migrate from the distal tip results in a radially symmetric proximal streak^6,7^. However, the proper head induction requires the secretion of WNT inhibitor DKK1 from the anterior axial endoderm rather than from the AVE^8^. Recent mouse gastrulation models (gastruloids), generated from mouse embryonic stem cells (mESCs), asymmetrically expressed the primitive streak protein BRACHYURY upon ligand stimulation in the absence of extraembryonic tissues ^9-11^. Furthermore, mESCs fused with mouse trophectoderm stem cells (TSCs) in a 3D matrix can self-organize into a cavitating embryo-like structure, which asymmetrically expresses BRACHYURY in the absence of added morphogens^12^.

To what extend these symmetry-breaking events are evolutionarily conserved and relevant to early human embryonic development is unknown. This question remains intriguing, as human embryos develop with a different architecture, and thus the shape and the juxtaposition of embryonic versus extraembryonic tissue is different than the mouse. Furthermore, the capacity of the embryonic tissue alone to break symmetry is unclear. Directly studying human gastrulation and symmetry braking remains extremely challenging, not only because of current ethical guidelines, but also because of the paucity of biological material^13^. Thus, synthetic models of human embryos provide the only alternative in addressing these questions. As mESC-based differentiation systems mimic aspects of *in vivo* gastrulation, conceivably human embryonic stem cells (hESCs) may also generate insights into human *in vivo* development^14^.

Here, we use hESCs to generate a 3D model of a human pre-gastrulation epiblast. By modeling the epiblast alone, we demonstrated its capacity to break symmetry and pattern upon a uniform application of the bone morphogenetic protein 4 (BMP4) and in the absence of extraembryonic tissues. BMP4 is one of the earliest morphogens necessary for the patterning of the mouse epiblast^15,16^ and it was shown to differentiate 2D micropatterned hESC colonies into organized germ layers driven by colony boundaries ^17,18^. Despite the fact that BMP4 could induce organized germ layers from patterned hESCs, no axial symmetry breaking was observed. Unlike in mouse embryo models generated from ESC/TSC aggregates, where the positioning of the TSCs created a bias for initiation of primitive streak genes, surprisingly we find that symmetry breaks spontaneously despite no apparent asymmetry in ligand application. Using a combination of genetic manipulations of the secreted inhibitors early embryonic signaling pathways and live-cell imaging, we demonstrate that model human epiblasts spontaneously polarize into regions expressing markers of the primitive streak and early ectoderm.

## RESULTS

### 3D human epiblast model

At 10 days of development, the human epiblast is an epithelial tissue lining the pro-amnionic cavity^19,20^. It was previously shown that human stem cells embedded in Matrigel spontaneously self-organize into spherical cavitating colonies^20-22^, however their precise fate and morphological properties remain poorly understood. We confirmed that Matrigel-embedded dissociated hESCs proliferated from single cells and formed cavitated 3D aggregates (Figure 1A). However, after three days in culture under pluripotency conditions, hESCs down-regulated NANOG, SOX2 and OCT4 and upregulated GATA6, GATA3, and BRACHYURY (BRA) indicating that they had differentiated, likely toward extraembryonic fates (Figure 1B, Figure S1A)^19,22,23^. Additionally, after three days, structural integrity was lost with the cells protruding into the surrounding matrix (Figure S1A) and it was blocked with LDN-193189, indicating that the process is BMP4 dependent (Figure 1B). As 3D hESC colonies in pure Matrigel differentiate despite the absence of added morphogens, we conclude that we cannot use this system to model a pre-gastrulating human epiblast.

**Figure 1:**
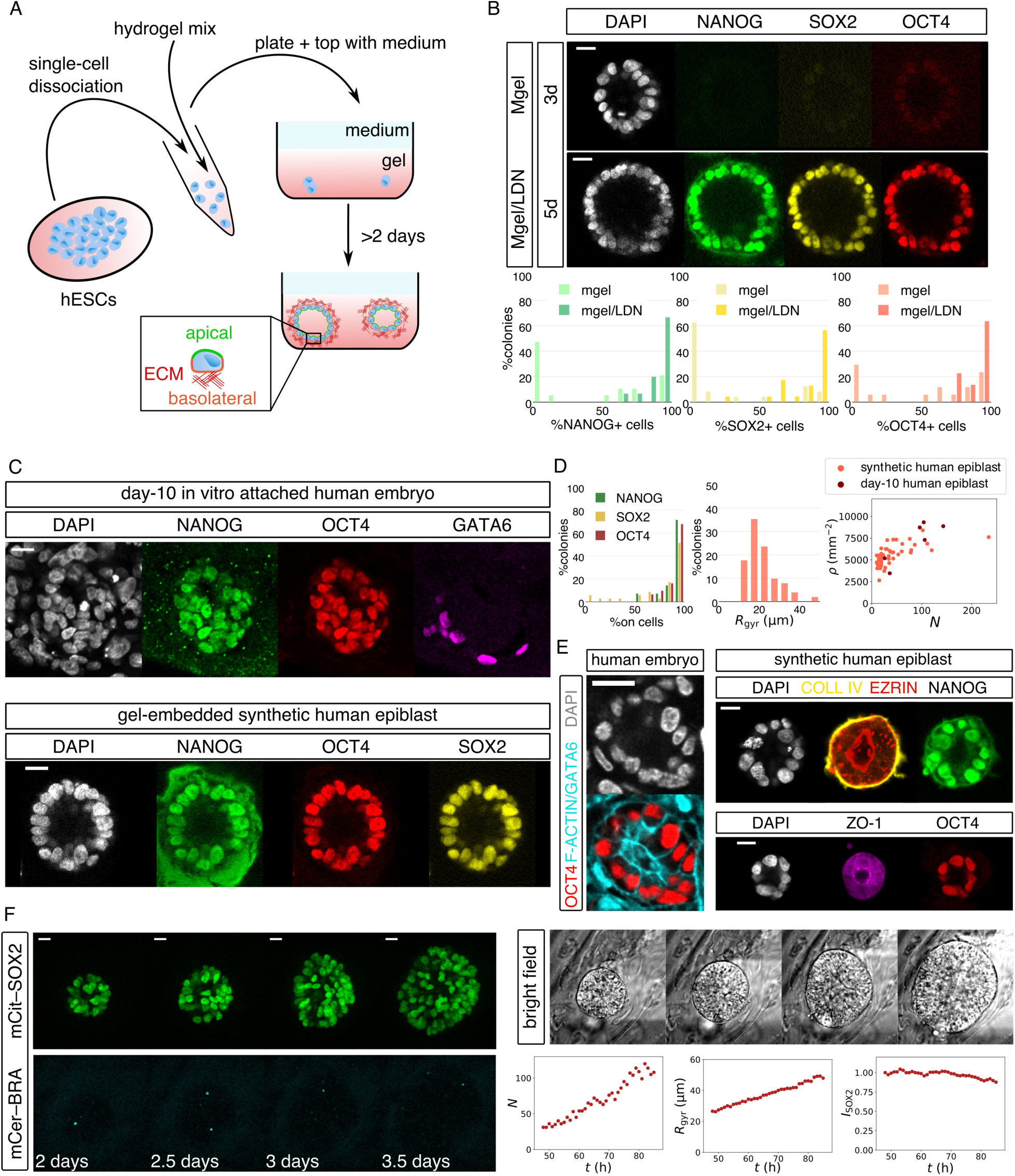
*In vitro* 3D model of a pregastrulation human epiblast. (A) Schematics of the experimental approach: dissociated hESCs are mixed into liquefied matrix (Matrigel or 4:1 (*V*/*V*) mix of hydrogel/Matrigel). (B) BMP-dependent spontaneous differentiation of cavitating hESC colonies in Matrigel. Shown are IF of pluripotency markers of a colony grown in Matrigel (Mgel) for three days and in Matrigel/LDN (Mgel/LDN) for five days. Plots quantify the pluripotency loss where each colony is binned based on the fraction of its cells expressing the pluripotency marker (*n* = 31 for Mgel; *n* = 28 for Mgel/LDN). (C) Comparing day-10 *in vitro* attached human embryo with a model epiblast grown in hydrogel/Matrigel for three days. In the embryo: NANOG+/OCT4+ cells mark epiblast, GATA6+ cells mark primitive endoderm. The remaining cells seen in DAPI are trophoblast and other extraembryonic tissues (see also Figure S1). (D) Pluripotency (left) and radius of gyration, *R*_gyr_, distribution (center) of a model human epiblast grown in hydrogel/Matrigel. The average *R*_gyr_ of the epiblast in attached human embryo was measured 29.7±2.2 μm (mean±s.e.m.; n = 6). Cell surface density, ρ, (right) *vs*. cell number, *N*, for 10-day human epiblast (*n* = 6) and synthetic epiblast (*n* = 51, data pooled from colonies grown over three or four days). (E) Apicobasal polarity of a day-10 and model human epiblast, showing IF staining of basal markers COLLAGEN IV (COLL IV) and apical markers F-ACTIN, EZRIN, and ZO-1 (human embryo: *n* = 4; synthetic epiblast: top row, *n* = 5, bottom row, *n* = 7). (F) Live-cell imaging of a synthetic epiblast under pluripotency conditions for ∼36 h starting on day two of culture. Each snapshot is a maximum intensity projection of *z*-slices. Plots quantify cell number, *N, R*_gyr_, and normalized SOX2 intensity, *I*_SOX2_, of the colony shown in micrographs. Shown is a representative example out of the *n* = 39 colonies that remained SOX2+ (out of the *n* = 46 total imaged colonies). All scale bars, 20 μm. Human embryo data was generated in our previous work ^19^. See more examples in Figure S2 and whole-population statistics in Figure S3.

To improve culture conditions, we used a polymeric hydrogel as the 3D matrix^24,25^ supplemented with Matrigel to provide a source of the basement membrane (Figure 1A). We cultured hESCs for two or three days to yield a wider size distribution. Under these conditions, the 3D hESC colonies displayed similar size range, morphology, molecular markers, and tissue polarity that matched our *in vitro* attached day-10 human embryos^19^. Namely, cells in both the *in vitro* attached embryo epiblasts and in our 3D model formed a quasi-spherical shell that surrounded a cavity, uniformly expressing NANOG and OCT4 (Figure 1C and S1B, C). Unlike in pure Matrigel, under these conditions cells expressed NANOG, SOX2, and OCT4 after four days in culture (Figures 1D and S1C). The average radius of ∼30 μm of the human epiblast together with the surface cell density compared very well with the model epiblasts (Figure 1D). Furthermore, they showed clear apicobasal polarity, with the outer edge lined by a network of COLLAGEN IV (COLL IV), one of key components of the basement membrane, and the inner side enriched with an actin–membrane linker protein EZRIN (Figures 1E and S1D). In addition, ZO-1, a protein localized at tight junctions at the apical side showed a pronounced expression at the cell-cell contacts on the interior of the model epiblast (Figures 1E and S1D). The *in vitro* attached embryo epiblast at day 10 is equally polarized seen by actin enrichment lining the pro-amniotic cavity (Figure 1E)^19,20^.

To monitor the dynamics of epiblast self-organization and the expression of key transcription factors, we recently created a fluorescent reporter of germ layers by tagging the endogenous loci for the key transcription factors using CRISPR/Cas9 into our hESC line (RUES2-GLR)^26^. This line expresses mCitrine-SOX2 (mCit-SOX2), active in pluripotency and anterior epiblast, mCerulean-BRA (mCer-BRA) expressed at gastrulation in the primitive streak and mesoderm, and tdTomato-SOX17 (tdTom-SOX17) expressed at gastrulation within endoderm and other tissues (Figure S2A). Starting 48h after seeding, live-cell imaging of RUES2-GLR model epiblasts under pluripotency condition showed a continuous proliferation over the course of at least 36h (Figures 1F and S2B, Movie S1), with 85% colonies remaining SOX2+/BRA-(*n* = 46) by the end of 3.5 days supporting our immunofluorescence (IF) data (Figures 1F, S2B, S3A, and S3B). In summary, based on morphological features, transcription factor and surface protein expression profiles, we use our 3D experimental system as a minimal model of a pre-gastrulation human epiblast.

### BMP4 concentration-dependent patterning of a model human epiblast

To test if the epiblast alone can break the AP symmetry in the absence of extraembryonic tissues and asymmetric ligand presentation, we, first, cultured hESCs in 3D for two or three days, to get a range of sizes, then applied BMP4 uniformly to the medium (Figure 2A). As the effective concentration of secreted BMP4 in a mammalian embryo is unknown, we tested concentrations from 0.1 ng/mL to 10 ng/mL. We checked the epithelial 3D colonies for the expression of known early gastrulation markers by imaging the end-point IF stains and by live-cell imaging of the RUES2-GLR line. Based on the presence of SOX2+ and BRA+ cells, we found three distinct populations of colonies: (i) pure SOX2+, (ii) colonies with SOX2+ and BRA+ cells, and (iii) pure SOX2− colonies. The fraction of each population within a system depended on BMP4 concentration (Figure 2B). At low BMP4 (0.1 ng/mL), the majority of colonies remained SOX2+ (Figure 2B). As live-cell imaging established that SOX2 levels remained constant over time (Figures S3B and S3C, Movie S2), and that end-point SOX2+ colonies were also OCT4+ (Figure S3D), we conclude that this population remains pluripotent. Thus, 0.1 ng/mL is not sufficient for differentiation. At high BMP4 (5 or 10 ng/mL) SOX2 declined uniformly and rapidly, with no obvious signs of asymmetry (Figures S3B and S3E, Movie S3). The majority of colonies expressed GATA6 and GATA3, with a subset also expressing BRA (Figure S3F). This molecular signature parallels observations made with high BMP4 concentration inducing extraembryonic cell types^27^. Finally, stimulating with the intermediate concentration of BMP4 (1 ng/mL) results in a mix of all three colony types, with approximately 50% of colonies yielding both SOX2+ and BRA+ cells (Figure 2B) providing a 3D platform to study axial patterning of the human epiblast model.

**Figure 2:**
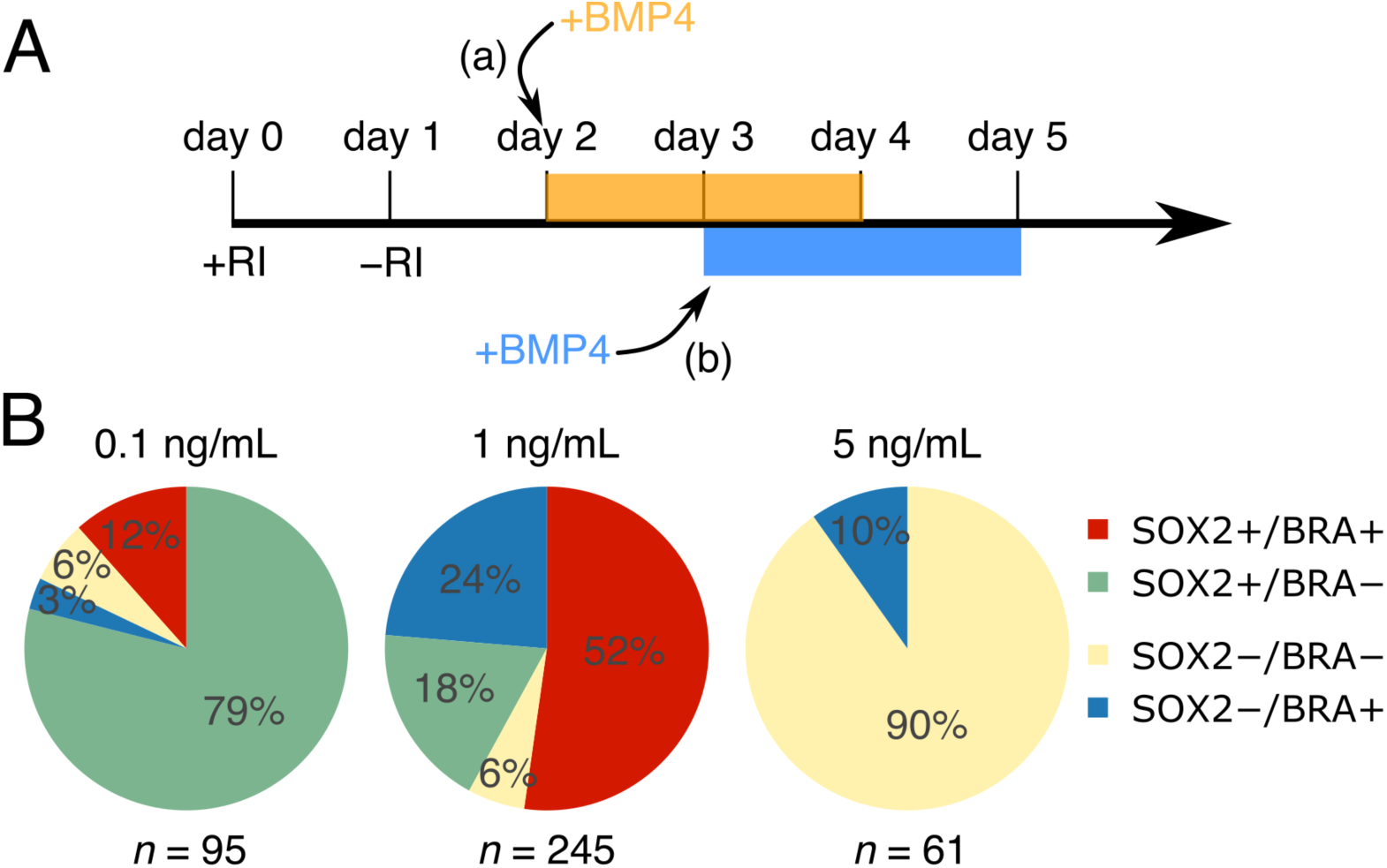
Concentration-dependent patterning of a model human epiblast by BMP4. (A) Schematics of the experimental protocol. BMP4 is added to the medium for two days either at day two or three of growth under pluripotency conditions (to ensure broader size distribution). RI, ROCK inhibitor. (B) Percentage fates of colonies under various BMP4 levels after two days. Sample size under each pie chart represents number of imaged colonies from combined end-point IF staining and live-cell imaging using the RUES2-GLR line (see Figure S3A for break-down). The composition of majority populations at 0.1 ng/mL and 5–10 ng/mL BMP4 is discussed in Figure S3. The 1-ng/mL populations are discussed in subsequent sections.

### Breaking the AP symmetry by intermediate BMP4 concentrations

In the colonies containing both SOX2+ and BRA+ cells, at 1 ng/mL BMP4, strikingly, the expression of the two cell types was typically asymmetric and spatially separated (Figure 3A). Spontaneous symmetry breaking was confirmed in seven independent experiments probing BRA/SOX2 expression as well as molecular markers of EMT, together with live-cell imaging. The influence of the particular synthetic hydrogel was ruled out by showing that the asymmetry occurred in chemically distinct hydrogels (Figure S4A).

**Figure 3:**
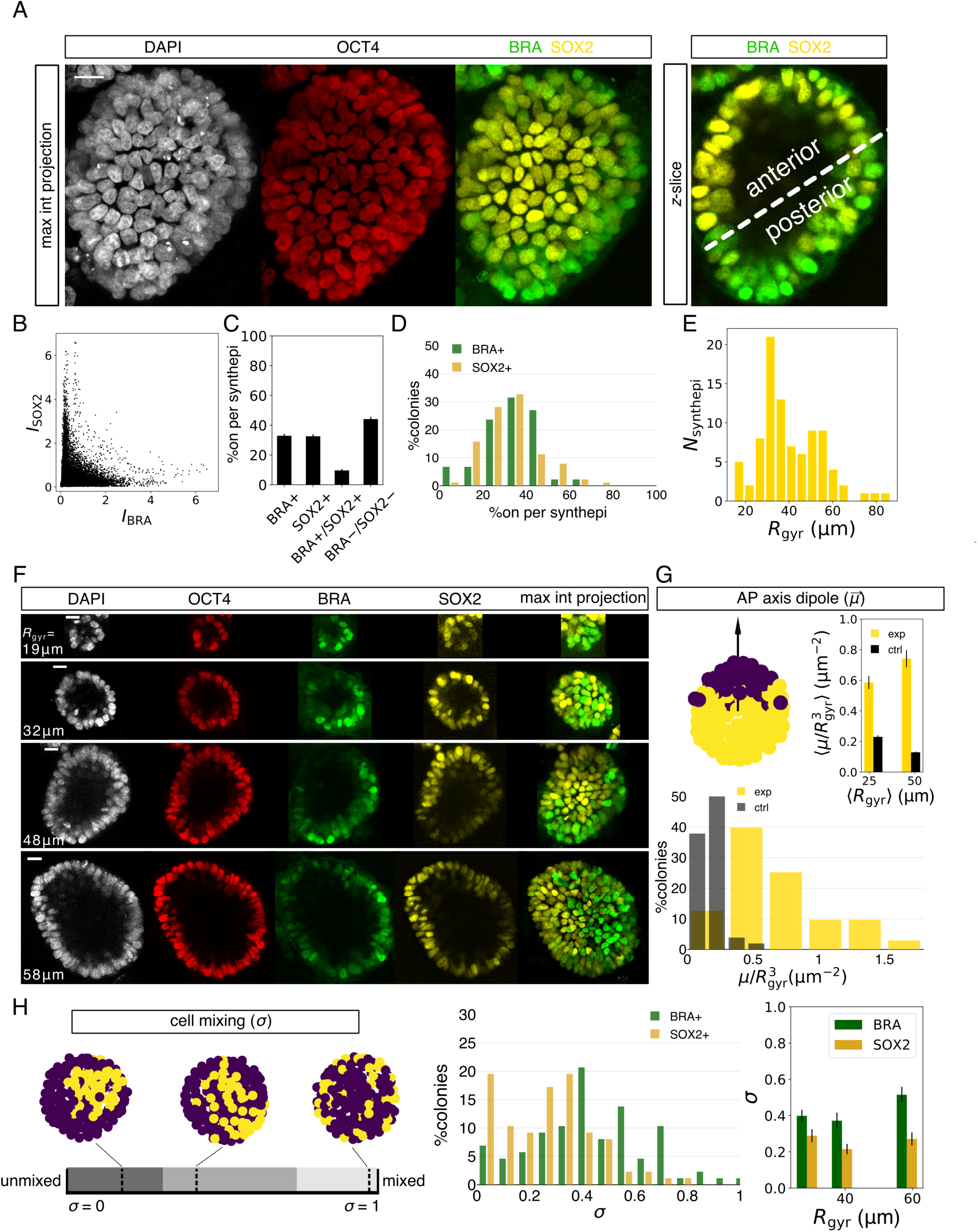
Breaking the AP symmetry in a model human epiblast. (A) Patterned model epiblast with asymmetrically expressed primitive streak marker BRA and the anterior marker SOX2 after two days in 1 ng/mL of BMP4. (B–E) Quantification of BRA/SOX2-expressing model epiblasts (*n* = 87). (B) SOX2 *vs.* BRA IF intensity of all cells from all BRA+/SOX2+ epiblasts. SOX2 intensity was rescaled by the ratio of population means of BRA and SOX2 intensities. (C) Mean fraction of cells expressing BRA, SOX2, both, or neither marker per model epiblast. Positive BRA or SOX2 signal is decided based on Otsu thresholding on individual channels. (D) Percolony distribution of cells expressing BRA or SOX2+ in BRA+/SOX2+ model epiblasts. (E) Size distribution of patterned model epiblasts. (F) Examples of BRA+/SOX+ model epiblasts across a wide size range, with *R*_gyr_ indicated on individual images. (G) AP dipole moment, 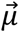. Top left: illustration of the AP vector on an example, with color-coding representing two populations (purple: BRA, yellow: SOX2). Top right: mean volume-normalized magnitude of AP dipole, ⟨μ/*R*_gyr_^3^⟩ of the population (exp) compared with a randomized control (ctrl). Data is binned into two size groups with a similar sample number (center between yellow and black bar is mean *R*_gyr_ of population). There is significant difference between the experimental *vs*. the scrambled colony for both bins (*p* < 0.0001) and only a moderate difference between experimental dipole values of two size bins (*p* = 0.03), assessed with the Mann-Whitney U test. Bottom: distribution of μ/*R*_gyr_^3^ from experiments (exp) and in controls (ctrl). (I) Cell mixing, σ, normalized by the maximum and minimum σ from 10,000 random permutations of cells. *Left*: an illustration of σ. It broadly captures three general features; low σ: compact roughly circular domains, intermediate σ: compact streak-like domains, and high σ: salt-and-pepper pattern. The compact domains in the illustration are generated from real examples, whereas the fully mixed epiblast example is simulated. *Center*: distribution of measured σ for the BRA and for the SOX2 population. *Right*: mean σ binned into three size groups of similar sample number and calculated separately for BRA+ and SOX2+ domains. Statistical significance for difference between σ(BRA) and σ(SOX2) from left to right: *p* = 0.03, 0.004, 0.0003, assessed with the two-sided Mann-Whitney U test. There is no statistical difference in σ for different size groups: *p* = 0.04 (BRA) and 0.4 (SOX2), assessed with the Kruskal-Wallis H test. All scale bars, 20 μm. Error bars in all panels are s.e.m.

The axial separation of individual cells within a colony into SOX2+ and BRA+ domains was not previously observed in human gastruloids^17,28,29^, which displayed only a proximal-distal axis, but instead, it mirrors gastrulation in the mammalian embryo where BRA defines the primitive streak and SOX2 marks the anterior epiblast. Of note, there was no appreciable expression of GATA3 in the symmetry-broken colonies, suggesting the absence of extraembryonic fates (Figure S4C) ^30^.

We quantified symmetry breaking in differentiated 3D colonies that contained both BRA+ and SOX2+ cells. First, the expression levels of individual markers showed that BRA and SOX2 expression was largely mutually exclusive (Figure 3B). Next, we partitioned the cells into expressing or nonexpressing, based on SOX2 or BRA IF intensity (see Figure S4B for an example of a segmented epiblast). ∼60% of cells per model epiblast were uniquely BRA+ or SOX2+, with roughly equal fraction of either marker (Figure 3C), and the distribution of colonies by composition peaked around an equal mixture (Figure 3D).

We observed symmetry breaking across the entire population of colonies regardless of size, in the range *R*_gyr_ = 15–80 μm (Figures 3E and 3F). There was, however, a difference in the morphology of the presumptive primitive streak among the population, with the BRA domain more spread in some colonies than in others (compare examples in Figures 3A and 3F). Given these differences, we devised two symmetry-breaking metrics. The first, AP dipole 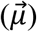, is a vector measuring the spatial segregation between the BRA and the SOX2 domains and is weighted by single-cell IF intensities (Figure 3G). As an analogy, this vector best distinguishes the distribution of two colors painting a sphere. Its value is zero if the colors are uniformly mixed, and it is at its maximum if the colors are segregated into separate hemispheres. In our system, 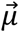 is aligned with the AP axis and its magnitude, *μ*, increases with the degree of symmetry breaking. By scrambling the IF intensities over the measured nuclear positions and repeating the calculation for the scrambled epiblast, we show that the measured 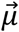 are very far from random, therefore quantitatively confirming symmetry breaking in patterned 3D colonies (Figure 3G). Furthermore, we observed that larger colonies polarize at only moderately higher dipole than the smaller colonies (Figures 3F, 3G, and S4C).

The second symmetry breaking parameter, cell mixing, *σ*, quantifies the compactness of the two domains. It is calculated by counting the number of nearest neighbor cell pairs expressing the same marker. Low σ values (<∼30%, dark stripe in Figure 3H) indicate an unmixed and a compact domain, intermediate values (∼30–75%, grey region) indicate an unmixed but more elongated pattern, whereas high values (>∼75%, light grey region) indicate perfect mixing, i.e. a salt-and-pepper pattern. Measuring *σ* across the entire BRA+/SOX2+ population, independently of μ, confirmed that the patterned model epiblasts were highly polarized, and again, irrespective of colony radius (Figure 3H). Interestingly, however, the mean *σ* of 20% for SOX2 compared to 40% for BRA indicates that SOX2 domain is more compact, while the BRA domain, although still unmixed, displays a more elongated pattern, i.e., more perimeter per area (Figure 3H).

### Epithelial-to-mesenchymal transition (EMT) of a model epiblast

To show that the symmetry-broken population is undergoing EMT and exhibiting molecular signatures consistent with the primitive streak, we examined a canonical marker SNAIL^31^ (Figure 4A, 4B and 4C). SNAIL was expressed and single-cell analysis of colonies expressing SNAIL and SOX2 confirmed quantitative symmetry breaking, marked by a shift in *μ* from the control and, independently, by low values of *σ* (Figure 4B). In the SOX2+ region, NANOG expression was very weak or absent, suggesting these cells are undergoing differentiation and thus are no longer pluripotent epiblast hESCs (Figure 4A).

**Figure 4:**
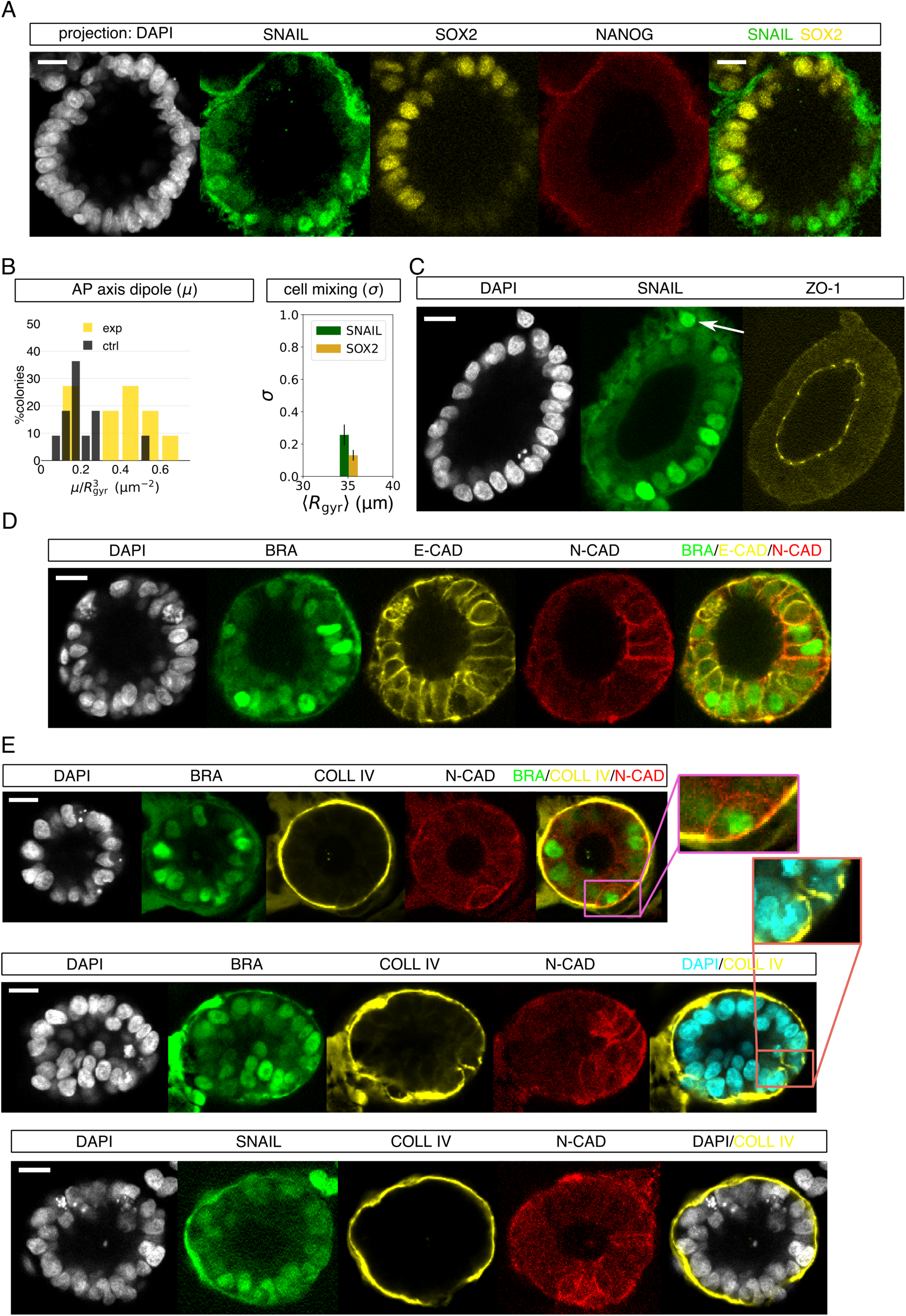
Signs of the primitive streak and EMT. (A) Primitive streak marker SNAIL is expressed asymmetrically to SOX2 (*n* = 11 for the SNAIL/SOX2 combination, of which 9 were stained for SNAIL/SOX2/NANOG). (B) Symmetry-breaking analysis on SNAIL+/SOX2+ patterned epiblasts (*n* = 11). SNAIL+ cells were determined as >0.9x Otsu threshold (see Methods). There was statistical difference between the means of the measured and the randomized AP dipole (p = 0.02), assessed with the two-sided Mann-Whitney U test, and no statistical difference between σ(SNAIL) and σ(SOX2) (*p* = 0.2), assessed with the Mann-Whitney U test. (C) Tight-junction marker ZO-1 is still expressed apically (*n* = 8 for SNAIL/ZO-1 combination). Arrow points to escaped cell not showing ZO-1. (D) Molecular signature of EMT: downregulation of adherent junction protein E-CADHERIN (E-CAD) and the upregulation of N-CADHERIN (N-CAD) in the BRA+ region (*n* = 3 for BRA/E-CAD/N-CAD combination). (E) BRA+/SNAIL+/N-CAD+ regions show degradation of basement membrane component COL-LAGEN IV (COLL IV). *Top* example shows the thinning of COLL IV in the N-CAD+ region, *center* example shows complete degradation, magnified in the inset (*n* = 6 for BRA/COLL IV/N-CAD combination). *Bottom*: COLL IV breakdown is also seen in the asymmetrically expressed SNAIL and N-CAD region (*n* = 6 for SNAIL/COLL IV/N-CAD combination).

The tight junction protein ZO-1 was still expressed at the apical side in cells that were part of the epithelium in SNAIL+ domains, although not in the nearby escaped SNAIL+ cells (Figure 4C). Moreover, the expression of cadherins showed a typical molecular signature of EMT, with a downregulation of E-CADHERIN^32^ and an upregulation of N-CADHERIN in the BRA-rich region (Figure 4D).

Strikingly, among the symmetry-broken colonies, there were clear signs of the break-down of the basement membrane in the BRA+ or SNAIL+ domain, ranging from a diminished COLL IV expression (Figure 4E, *top*) to a complete disintegration of the collagen network (Figure 4E *center* and *bottom*). COLL IV breakdown overlapped with N-CADHERIN expression, confirming EMT^33^ (Figure 4E). Collagen breakdown in the primitive streak is a prominent sign of gastrulation in the mouse embryo^34^. Taken together with our previous observations human micropatterned gastruloids and molecular signatures from the mouse, the asymmetric expression of BRA, SNAIL, and N-CADHERIN and the breakdown of the basement membrane demonstrate that our 3D model system mirrors the initial steps of AP axis specification^14,17^.

### Dynamics of symmetry breaking

We monitored the dynamics of symmetry breaking by time-lapse imaging of model epiblasts composed of the RUES2-GLR line. At intermediate BMP4 concentrations (1 ng/mL) half the colonies asymmetrically expressed BRA/SOX2, quantitatively comparable to IF measurements (Figures 5A, S3A, Movies S4-S6). In a fraction of the population the SOX2 level remains unchanged, suggesting they remain pluripotent, again consistent with IF measurements (Figure S5A,B). Although the signal-to-noise ratio is much lower in live-cell imaging due to the gel-embedding, low endogenous protein expression, and lowered imaging power to prevent phototoxicity, time-lapse clearly showed that, in the BRA+/SOX2+ population, SOX2 expression declines and BRA expression begins in one spot and then expands (Figure 5A). Consistently, the quantification shows that concurrent with the BRA activation at 20–25h post BMP4 induction, the AP dipole sharply rises (Figure 5B). Therefore, the two markers do not stochastically turn on in a salt-and-pepper pattern then sort out; rather, BRA activation is a result of signaling asymmetry.

**Figure 5:**
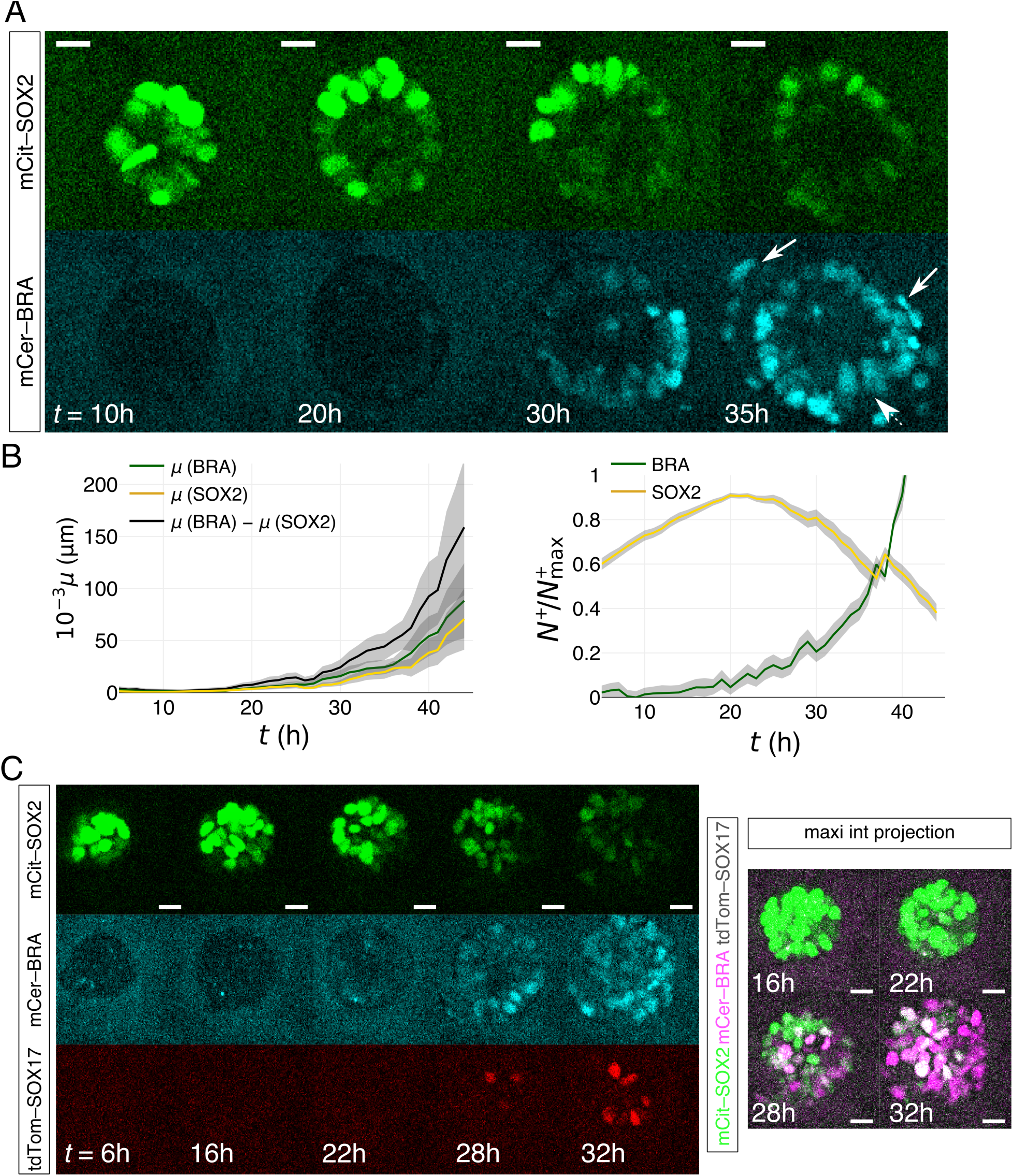
Dynamics of AP symmetry breaking, EMT, and SOX17 activation. (A) Live-cell imaging of a RUES2-GLR model epiblast. Time is measured relative to addition of 1 ng/mL BMP4. The arrow points to cells undergoing EMT. (B) Symmetry-breaking quantification of BRA+/SOX2+ colonies (*n* = 34 BRA+/SOX2+ colonies). *Left:* volume-uncorrected mean *μ* measurement from all BRA+/SOX2+ movies, broken down into the individual *μ* contributions from BRA+ and SOX2+ voxels. *Right:* SOX2 *vs.* BRA expression in time, measured as mean number of BRA+ or SOX2+ voxels in the movie, *N*^+^, corrected by the maximum number of expressing voxels for each movie, *N*^+^_max_. The initial increase in SOX2 is due to the growth of the 3D colony. Shadow in both plots represents s.e.m. (C) Live-cell imaging of RUES2-GLR in all three channels, showing the activation of SOX17 (observed in 10 out of 11 BRA+/SOX2+ synthetic epiblasts, *n* = 19 total imaged epiblasts in all three channels). Confocal slices among different channels are different, but same for each channel. All scale bars, 20 μm.

After establishing the BRA/SOX2 axis, EMT was often observed in BRA+ cells escaping the epithelial tissue which sometimes underwent involution, hence suggesting morphogenetic movements, a hallmark of gastrulation (Figure 5A, Movie S5). Finally, despite the absence of the enveloping extraembryonic tissue, which would limit the BRA+ expansion and its escape into the medium, SOX17 expression was nevertheless observed following BRA activation. We thus captured early specification events as expected in gastrulation (Figure 5C and S5C, Movie S6).

### Molecular mechanism of symmetry breaking

In light of our recent finding that the BMP4 receptors are localized at the basolateral side of hESCs, we expect all cells to have equal initial access to BMP4^29^. To confirm the uniformity of the initial BMP4 response we monitored the dynamics of SMAD1 translocation into the nucleus by live imaging model epiblasts composed of RUES2 line with fluorescently tagged SMAD1^35^. At both the intermediate (1 ng/mL) and high (10 ng/mL) BMP4 concentration, SMAD1 nuclear shuttling was uniform (Figure S6, Movie S6). At the intermediate concentration, only half the colonies showed SMAD1 signal at 8h and beyond, comparable to the population fraction that displayed polarized BRA/SOX2 expression (Figure S6). Thus failure to differentiate may be a reflection of the initial BMP4 response.

Recently, we have elucidated the BMP4:WNT:NODAL signaling cascade in 2D micro-patterned hESCs, which relied on the BMP4 inhibitor NOGGIN to induce radially organized germ layers^29^. Furthermore, we identified a variety of secreted WNT inhibitors produced upon WNT3A stimulation of 2D micropatterned colonies^26^. Thus, we sought to identify the key components involved in the AP induction in our model epiblasts. To get an adequate yield of cells for gene expression analysis, instead of using 3D colonies, we grew hESCs on transwell filters. These have been shown to promote the formation of a large, contiguous, and uniform epiblast-like epithelium^29^. Moreover, transwell filters allow basal application of BMP4 uniformly to all cells in a manner that is equivalent to the 3D system. After titrating BMP4 levels, we found that a 10-ng/mL dose of BMP4 in the bottom compartment of the transwell filter for 24h results in segregated BRA+ domains intermixed among the SOX2+ cells (Figure 6A). As we previously found that the initial response to high BMP4 is uniform, this result is again an instance of spontaneous symmetry breaking reinforcing the similarly of the filter system to the 3D epiblasts^29^.

**Figure 6:**
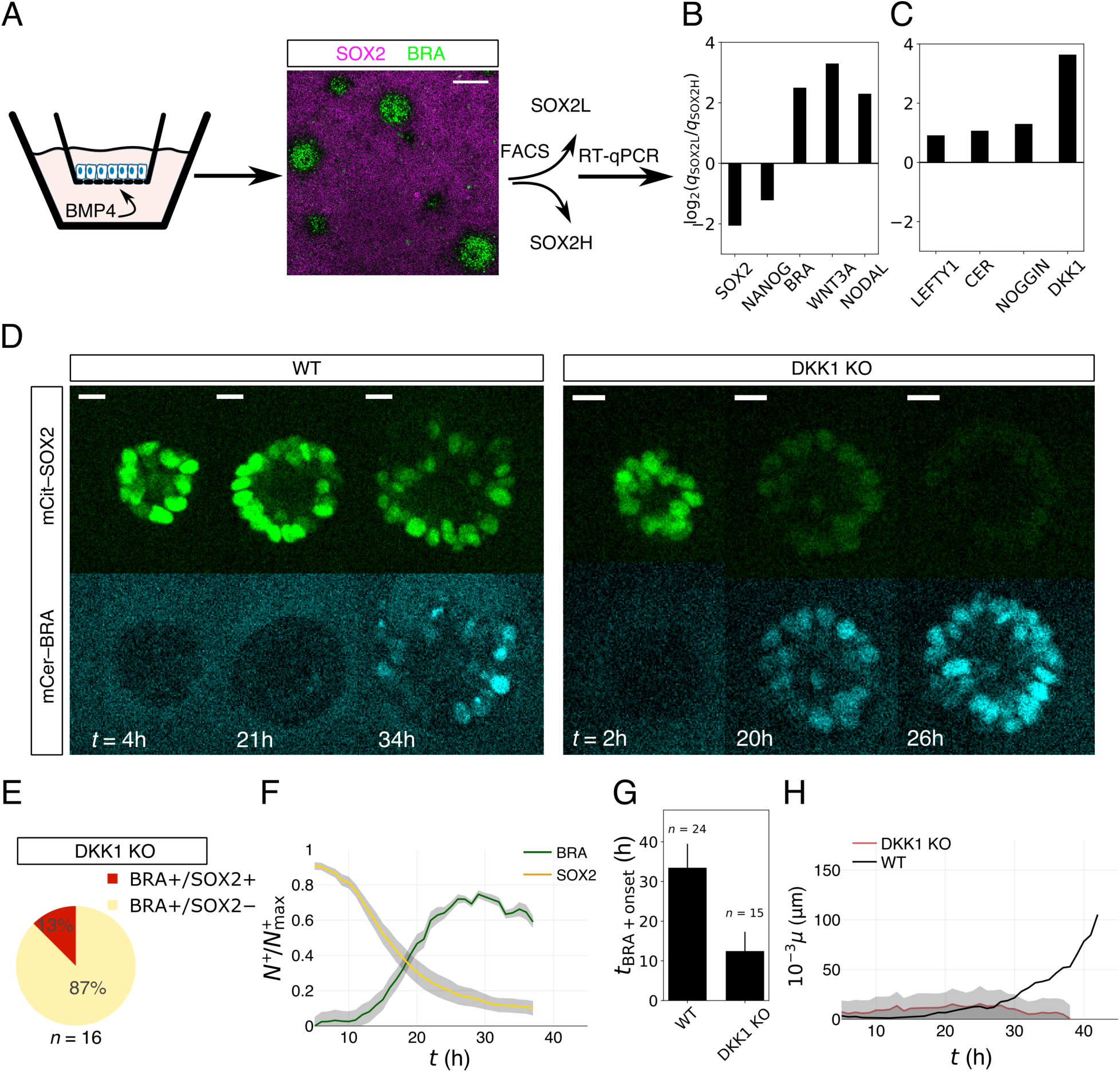
The molecular mechanism of symmetry breaking in the model epiblast. (A) Stimulating RUES2-GLR 2D model epiblast grown on transwell filters with 10 ng/mL BMP4 from the basal side for 24h yields regionalized BRA+ and SOX2+ patterns. Tissue is dissociated then FACS sorted based on mCit-SOX2 expression into high (SOX2H) and low (SOX2L)-expressing cells. Example patterned tissue shown is made from RUES2 and IF-stained for clarity (see more examples in Figure S6). Scale bar, 200 μm. (B-C) RT-qPCR of sorted cells. Shown are log2 of the expression ratio between cells in the SOX2H and SOX2L regions. (D) Live-cell imaging of RUES2-GLR and RUES2-GLR DKK1 KO model epiblasts stimulated with 1 ng/mL BMP4. Ze-ro-time represents ligand application. Scale bar, 20 μm. (E) Patterning statistics of all live-cell imaged DKK1 KO model epiblasts. The majority population (BRA+/SOX2-) is the one that uniformly deactivates SOX2 and activates BRA. (F) Protein expression analysis, measured as BRA+ or SOX2+ voxel number, *N*^+^, normalized by the maximum number of expressing voxels of the individual colony, *N*^+^_max._ Errors are s.e.m. (G) Average onset of BRA appearance after stimulation with 1 ng/mL BMP4 of RUES2-GLR (i.e., WT) and RUES2-GLR DKK1 KO model epiblasts, deduced from live-cell imaging. (H) Volume uncorrected μ measured from live-cell imaging of the RUES2-GLR DKK1 KO line (only analyzing the majority population), showing that no dipole is being induced during the experiment. The measurement is compared to the symmetry-broken population in the RUES2-GLR (i.e., WT; black line) reproduced from Figure 5B. Errors are s.e.m.

We then FACS-sorted the patterned 2D tissue into high- and low-SOX2-expressing cells then performed RT-qPCR on the two pools (Figure S7A). The SOX2-low region highly expressed *BRA, WNT3A*, and *NODAL* (Figure 6B), consistent with the primitive streak genes expressed in the mouse^16^. There was no appreciable difference in the expression of NODAL inhibitors *LEFTY1* and *CERBERUS* or the BMP inhibitor *NOGGIN* (Figure 6B,C). *DKK1* was also highly expressed in the BRA+/SOX2– region at 24h (Figure 6B,C). While the expression of the activator (Wnt) and its inhibitor (Dkk1) in the same tissue may seem surprising, it is consistent with their localization *in vivo* and theoretical expectations (see Discussion).

To confirm that WNT/DKK1 pair is key in symmetry breaking of model epiblasts, we knocked out DKK1 in our RUES-GLR line and conducted live-cell imaging upon adding 1 ng/mL BMP4. We find that in stark contrast to the WT that asymmetrically expressed the two markers, in >80% of colonies, RUES2-GLR DKK1 KO rapidly and uniformly lost SOX2 expression then uniformly activated BRA+ (Figure 6D,E, Movie S7). Shortly after BRA takes over the colony, most cells undergo EMT and the colonies disintegrate (Movie S7). Furthermore, stimulated RUES2-GLR DKK1 KO colonies displayed a precocious SOX2 loss/BRA activation, with BRA appearing on average ∼20h earlier than in the WT (Figures 6F,G). As a control, we confirmed that under pluripotency and the same imaging conditions, the majority of RUES2-GLR DKK1 KO model epiblasts remained SOX2+ (Figure S7B). Finally, the absence of symmetry breaking is also quantitatively demonstrated by a low dipole that remains nearly constant throughout imaging (Figure 6H). Adding the Wnt secretion inhibitor IWP2 along with BMP4 aborted the activation of BRA, confirming that symmetry breaking is WNT-dependent (Figure S7C).

As hESC grown on filters provide an opportunity to independently confirm key effectors in the symmetry breaking mechanism, we stimulated RUES2-DKK1 KO cells with 10 ng/mL BMP4 from the bottom compartment. Consistently with the 3D model, while the WT displayed patches of BRA expression, symmetry breaking was completely abolished on 2D DKK1 KO epithelia (Figure S7D).

To test the effect of inhibitors of the BMP4:Wnt:Nodal signaling cascade, we generated model epiblasts from RUES2 NOGGIN KO and from RUES2 LEFTY1/CERBERUS double KO^36^. NOGGIN is the predominant secreted BMP4 inhibitor active in 2D micropattern gastruloids^29^. NOGGIN KO model epiblasts stimulated with 1 ng/ml BMP4 almost entirely lost SOX2 expression (Figure S7E), suggesting the loss of anterior fate. The double LEFTY1/CERBERUS KO yielded symmetry broken epiblasts as defined by SOX2/BRA at similar frequencies to WT, suggesting that NODAL is not necessary in the initial axial patterning.

## Discussion

In recent years, considerable progress in mammalian and, particularly, human development has been made by recapitulating aspects of early embryogenesis *in vitro*^37^. The establishment of the AP axis in mammals, in contrast to lower vertebrates where sperm entry provides the initiating bias, appears to be stochastic. In the mouse, the initiation and the positioning of the primitive streak is thought to be under the direct control of Wnt from the posterior visceral endoderm and the Wnt and Nodal inhibitors secreted from the AVE^5,16,38,39^. However, the organizing capacity of the epiblast itself remains an open question. Our work establishes the minimal cellular and molecular components for observing the AP symmetry breaking in a human epiblast model. Our observation stands in contrast to prior work with synthetic mouse embryos, where an asymmetric source of BMP4 was required^12^. We show that, surprisingly, a human epiblast model can spontaneously break the AP symmetry and undergo EMT upon uniform stimulation with BMP4, and in the absence of extraembryonic tissues and maternal cues.

Live imaging showed symmetry breaking did not begin as a stochastic expression followed by cell sorting but rather coherently over the colony, suggesting that the epiblast responded to a signaling gradient. It is tempting to speculate that the lumen in the 3D model epiblasts plays important roles, such as concentrating the secreted inhibitors, and possibly explaining why symmetry breaking was not observed in 2D systems.

While our study identifies the molecular players involved in symmetry breaking, unraveling the precise biophysical mechanism is extremely challenging. What comes as a surprise is that both the activator and the inhibitor come from the epiblast and from the presumptive streak, unlike the prevailing picture in the mouse where the DKK1 is typically seen coming from the AVE. Interestingly, however, recent high-resolution RNA sequencing of the E7 mouse epiblast^40^, as well as earlier work in rabbit ^41^, showed that Dkk1 is also elevated in the primitive streak and not just in the AVE. The earliest evidence for Dkk1 secretion in primates comes from a recent experiment in the macaque embryo where Dkk1 was detected in the AVE before gastrulation, however there is no data on Dkk1 localization at gastrulation in primates^42^. The spatial overlap of morphogen activity with its inhibitor conforms to patterning mechanisms based on reaction-diffusion or Turing models^43^.

Our work provides first glimpse of AP symmetry breaking and axis specification that might occur in human embryos as seen in our 3D model epiblast. Our assay provides a foundation for future exploration of the influence of the mechanical environment and extraembryonic tissues in the early patterning, a key step in resolving the interrelation of morphogenesis and fate acquisition in human embryogenesis.

## METHODS

### Cell culture

RUES2 hESC line (NIHhESC-09-0013) was used and cultured in HUESM conditioned by mouse embryonic fibroblasts (MEF-CM) and supplemented with 20 ng/mL bFGF. Cells were grown at 37 °C and 5% CO_2_ on tissue culture dishes which were coated with Matrigel (BD Biosciences) at 40x (*V*:*V*) dilution. Cells were passaged every ∼five days as small colonies or as single cells. Medium was changed daily. Cells were regularly checked for mycoplasma infections and they were used for no more than ∼20 passages.

### CRISPR/Cas9-mediated genetically modified cell lines

All genetic modifications were conducted in the RUES2 background. All cell lines and reporters were previously created and validated for pluripotency and genetic integrity. The RUES2-GLR line (the germ layer reporter) was created and validated in ^26^. RUES2-GLR DKK1 KO, RUES2 DKK1 KO, and RUES2 LEFTY1 & CERBERUS1 KO were created and validated in ^36^. RUES2 NOGGIN KO was created and validated in ^29^. RUES2 SMAD1 reporter was created and validated in ^35^.

### *In vitro* cultured human embryos

Images of IF stained day-10 human embryos in Figures 1 and S1 were generated as part of previous work ^19^.

### 3D model of a human epiblast

The experiment was performed in a 35-mm plastic imaging dish (Ibidi). Cells were dissociated as single cells using Accutase (Stem Cell Technologies), spun at 300g for 3 min, and resuspended with MEF-CM + bFGF to 1–2×106 cells/mL. ∼30 μL of cell suspension was mixed with 30 μL Matrigel and 150 μL hydrogel on ice, in that order (see below for hydrogel chemistry). Gels were kept on ice the entire time and handled rapidly to prevent premature solidification. The components were thoroughly mixed, taking care not to introduce bubbles, then transferred to the bottom of the dish in a snake-like pattern, to fully cover the inner circle of the imaging dish. The gel-embedded cell mix was placed in the incubator for 5–10 min to solidify then topped with warm MEF-CM + bFGF + pen/strep, supplemented with 10 μM ROCK inhibitor Y27632 (RI) for ∼24 h (Figure 2A). In case the gel-mix did not adhere to the surface, medium change was done by spinning down the suspension at 150 g for 1 min.

We tested two chemically distinct hydrogels: PEG-based ^24^ (Mebiol, Cosmobio) and Fmoc-based ^25^ (Biogelx). Mebiol was prepared by diluting the lyophilized gel with 10 mL PBS+/+ (Sigma) overnight. Biogelx was prepared in aliquots, by diluting 21.5 μg of the powder with 5 mL water then vigorously vortexing for several minutes. Both liquefied hydrogels were kept at 4 °C. The quality of the hydrogels and the Matrigel deteriorated in the fridge, as judged by reduced cell survival when using the gel refrigerated for a long time. Mebiol and Biogelx were used within 2–4 weeks after the initial dilution. Biogelx required vigorous vortexing or sonication before each experiment. Matrigel was initially thawed in the fridge overnight then aliquoted and kept in the freezer. Each aliquot was then only re-thawed once and used within 2 weeks.

### Patterning of the human epiblast

To start the experiment, medium was changed (MEF-CM + bFGF + pen/strep) and supplemented with a desired concentration of BMP4 and/or other tested molecule. The dish was left unperturbed for 48h in the incubator or mounted into the incubation chamber on the microscope in case of live-cell imaging.

### IF staining and imaging of synthetic epiblasts

The experimental medium was collected, spun down at 150g for 1 min, then resuspended with 4% paraformaldehyde (dissolved in PBS+/+) and poured back into the experimental dish to fix the synthetic epiblasts for 30–60 min. After washing twice with PBS+/+, the experiment was blocked with 2% bovine serum albumin in 0.1% Tri-ton-X (Sigma), dissolved in PBS+/+ (block solution), for at least 30 min at 4 °C. This step lique-fies the gel and frees the synthetic epiblasts into the solution. Block was replaced with primary antibodies (see Table S1 for catalog numbers and dilutions), dissolved in block solution overnight. Primary antibodies were washed away three times with 0.1% Triton-X solution in PBS+/+, keeping the solution 15–30 min each wash. Then, secondary antibodies and DAPI diluted in block solution were added for 30 min. Finally, the synthetic epiblasts were washed twice with 0.1% Triton-X in PBS+/+ and kept in that medium at 4 °C. Most synthetic epiblasts sink and adhere to the bottom of the dish.

We list the antibodies and dilutions for the following antigens: BRACHYURY (R&D Systems AF-2085; 1:300), SOX2 (Cell Signaling 3579; 1:200), OCT4 (BD Biosciences 611203; 1:200), NANOG (R&D Systems AF-1997, 1:150), SNAIL (R&D Systems AF-3639; 1:150), COLLAGEN IV (Abcam Ab-6586; 1:200), EZRIN (Sigma-Aldrich E8897; 1:200), GATA3 (ThermoFisher, MA1-028; 1:100), GATA6 (Cell Signaling 5851S; 1:200), E-CADHERIN (Cell Signalling 3195; 1:200), N-CADHERIN (BioLegend 350802; 1:200), ZO-1 (Invitrogen 617300; 1:200).

Imaging of fixed samples was performed on Zeiss Inverted LSM 780 laser scanning confocal microscope with a 10x, 20x, or 40x dry objective. Live-cell imaging was performed with a Yokagawa/Olympus CellVoyager spinning disk microscope with a dedicated cell incubator (37 °C, 5% CO_2_) using a 20x dry objective. Images were acquired at a frequency of 1h^−1^ for no longer than 40h at laser powers and exposure times to optimize for minimum cell death.

### A note on preparation variability and reproducibility

Both tested hydrogels yielded symmetry-broken structures (Figure S4B) demonstrating the robustness of the phenomenon. Preparation quality of synthetic epiblasts was strongly dependent on how cells and gels were mixed. In case of poor cell disassociation, large clumps would form and resembled embryoid bodies. In the case of Mebiol, improper gel mixing would result in a patchy gel mix and therefore a coexistence of pluripotent synthetic epiblasts and differentiated colonies as part of the Matrigel patch. In general, nonstructured cell clusters such as loose mesenchymal cells or disintegrating structures were ignored. This issue, however, was almost completely absent when using Biogelx. Importantly, cell survival and thus the quality of the experiment deteriorated with refrigeration time of hydrated gels and of Matrigel.

### Patterning and imaging hESC on filters

Transwell polycarbonate membrane (Costar) inserts, 12 mm in diameter, were coated for 2 hr at 37 °C with 250 mL of 10 mg/mL laminin 521 (Biolamina) diluted in PBS+/+. The membrane was rinsed with PBS+/+. Cells were dissociated as single cells with Accutase and resuspended with MEF-CM + FGF +pen/strep + 10 μM RI. To each filter insert, 500 μL of the same medium containing 100,000 cells was carefully added atop the membrane, while ∼1.5 mL of the same medium (not containing the cells) was added to the bottom. RI was washed out the next day (from both filter compartments). To the bottom of the well, 48h after seeding, MEF-CM + FGF + pen/strep + 10 ng/mL BMP4 was added. Filters were fixed 24h later, stained as described above then imaged with a 20x objective.

### FACS sorting and qPCR of patterned hESC filters

100,000 RUES2-GLR cells were seeded on 6.5 mm polyester filters in MEF-CM + FGF + pen/strep + 10 μM RI (*n* = 15 replicates). Two hours after seeding, medium was replaced by MEF-CM + FGF + pen/strep. The next day, a confluent layer of pluripotent hESC formed and 10 ng/ml BMP4 was added to the basal side of ten filters, while five filters were left in pluripotency medium as control. After 24h, the cultures were dissociated as single cells with Accutase and samples associated with differentiation or control conditions were pooled together. By FACS sorting we defined SOX2 high and SOX2 low cells based on mCit-SOX2 expression (Figure S6A). Live-to-dead gating was determined based on forward scattering. On average, 70% of the cells were gated as live. To determine SOX2-high and SOX2-low populations, gating of the SOX2-high population was determined in the undifferentiated sample. This led, in the differentiated sample, to 67% of cells in the SOX2-high gate and 25.5% of the cells in the SOX2-low gate. Differential gene expression in these three populations was analyzed by qPCR using standard protocols.

### Analysis of IF stained colonies

First, it should be noted that the brightness of fluorescence images was adjusted to be visually representable in figures and was always done equally and linearly across the entire image. Confocal images of individual synthetic epiblasts were segmented based on DAPI staining. First, the background of individual images in all channels was subtracted using the rolling ball background subtraction algorithm in Fiji. Seeds for single-cell segmentation were generated by Ilastik, and used by an in-house watershed-based algorithm to generate the segmented images. Fluorescence intensities of individual cells were taken as median intensity, divided by median DAPI intensity of that cell. Based on visual inspection of colonies rendered in 3D, some distortion in *z*-axis was observed, likely contributed by the adhesion, centrifugation, and by light aberration, inherent to the dimensionality of our structures and the gel remnants after staining. No corrections were made, as it did not appreciably affect the normalized symmetry-breaking parameters.

Radius of gyration, *R*_gry_, was calculated as:

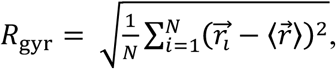

where *N* is the total number of cells, 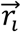 is the position of the centroid of cell *i* and 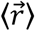 is the mean position of all centroids. Cell surface density, *ρ*, was defined as:

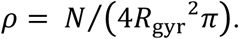

Borrowing from chemistry, AP dipole was calculated equivalently to the electric dipole of a molecule, where a segmented cell was equivalent to an atom and its median IF intensity to an effective charge. First, fluorescence intensities for each colony were rescaled to the maximum per-cell median intensity in each channel. This step was done to ensure equal scaling across the population, as the intensities from IF stains generally cannot be compared between experiments and they are also affected by the height of the imaged colony in the chamber. Then, a dipole for BRA and SOX2 were separately calculated as:

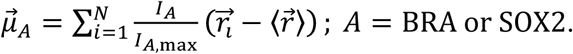

The total dipole, 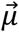, and its magnitude, *μ*, are then calculated as:

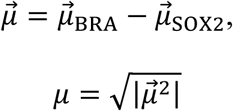

and normalized by the volume dimensions, *μ*/*R*_gyr_^3^, to remove the expected dependence on colony size in the population. In theory, any nonzero value indicates symmetry breaking, however noise can give rise to small *μ*. To ensure the measured *μ* quantitatively demonstrates symmetry breaking, for each patterned synthetic epiblast, we permutated the measured SOX2 and BRA intensities among segmented cells and calculated the mean *μ* from 1000 permutations.

Cell mixing parameter, *σ*, was inspired by edge energy of immiscible liquids – the smaller the boundary between the two liquids, the lower the mixing. In our case, *σ* is a fraction of non-like neighbors of a cell, averaged for all cells of a colony. To extract the network of neighbors, centroids of the segmented image were first surface meshed with a 3D Voronoi tessellation, which produced a list of “edges”, i.e. cell–cell contacts. Then, each cell was determined to be BRA+ and/or SOX2+ by Otsu thresholding of individual colonies in each channel separately (in Figure 4, Otsu threshold for SNAIL was adjusted by a factor of 0.9 judged by visual inspection). Non-normalized cell mixing, 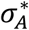, was calculated as:

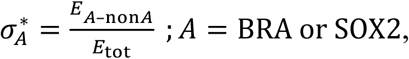

where *E*_*A*-non*A*_ is the number of non-like edges (in the case of BRA, those connecting a BRA+ and a BRA-cell), and E_tot_ is the total number of edges. The more cells are mixed, the higher σ will be, however the actual maximum and minimum depend on the number of BRA+ *vs* SOX2+ cells. Therefore, for each colony, the idealized minimum and maximum are calculated by permutating BRA+ and SOX2+ among the segmented cells. The normalized σ, which by definition runs from 0 to 1, as reported in the Main text is then calculated as:

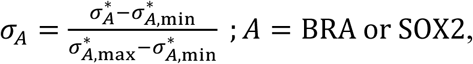

where 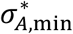 and 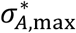 are, respectively, the minimum and maximum values among 10,000 permutations.

### Analysis of live-cell imaging

Note, brightness was adjusted for representation and it was done linearly and equally across the entire image. Since the colonies in 3D are suspended above the surface, their fluorescence signal will be different from example to example and should therefore not be compared in terms of absolute signal. Plots reported in Figures 1F, S2B, and S2C were segmented based on SOX2 expression. Under induction conditions, however, due to the low endogenous expression of BRA and therefore a very weak signal, the analysis was done on voxel intensities. The image was binarized into BRA and SOX2-expressing voxels by Otsu thresholding. Given that the synthetic epiblasts were live imaged while still embedded in the gel, at various distances from the surface, and overall a lower signal-to-noise ratio compared to IF, the thresholding had to be adjusted for each colony due to different absolute signal intensities among colonies. The mean intensity was rescaled so that the first snapshot equals 1 for SOX2 and 0 for BRA for all, but plot in Figure S3B (right) where the first snapshot in SOX2 was set to 1 and the minimum to 0 for clarity. AP dipole was calculated as above accounting only for BRA i.e. SOX2 positive voxels which were not intensity-weighted. The measurement for each colony was done up to EMT, as escaping cells compromise *μ*. Movies with very low signal to noise ratio and, in a few cases, where non-related cells appeared in the proximity were discarded due to uncertainty. Blemishes in imaging, such as coming from autofluorescence of protein aggregates in the region of interest were not specifically eliminated; therefore, our measured effect might be underestimated for some cases. Live-cell imaging was not analyzed past 40h.

### Summary of data statistics

The number of colonies for each data analysis or representative image is listed in each figure caption (as *n*). The statistics in Figure 2 was generated from BMP4-differentiated populations where cavitating epithelial colonies were imaged regardless of protein expression (pooled from IF stains and live-cell imaging) and these experiments were done in Mebiol gel. Additional data for quantification in the remaining figures was collected where only BRA+/SOX2+ (or equivalent, e.g. SNAIL+/SOX2+) colonies were considered and imaged at higher resolution and these were conducted either in Mebiol or Biogelx gel.

### Code availability

Code used for cell segmentation in this work is available from the corresponding author upon reasonable request.

### Data availability

The data that support the findings of this study are available from the corresponding author upon reasonable request.

## ACKWNOLEDGEMENTS

We are thankful to all the members of the Brivanlou and Siggia labs for helpful discussions. M.S. is a Junior Fellow of the Simons Society of Fellows. We were supported by NIH grants R01 HD080699 and R01 GM101653 to A.H.B. and E.D.S. Imaging was performed at The Rockefeller University Bio-Imaging Resource Center. We thank Sophie Morgani (MSKCC) and the members of the Siggia and Brivanlou laboratories for helpful discussions and the critical reading of the manuscript.

